# let-7i-5p regulates osteogenic differentiation of bone marrow mesenchymal stem cells induced by psoralen by targeting CKIP-1

**DOI:** 10.1101/2024.03.20.585715

**Authors:** Ming Chen, Song Zhou, Sheng Huang, Hao Luo, Xuelong Zhang, Gangan Liu, Huanan Li, Bo Liu, Dongmei Yan

## Abstract

Previous studies have shown that psoralen can treat osteoporosis by inducing osteogenic differentiation of bone marrow mesenchymal stem cells (BMSCs). CKIP-1 is an inhibitor of bone formation and plays a negative regulatory role in the osteogenic differentiation of BMSCs. MicroRNA (miRNA) is a kind of small single-stranded non-coding RNA, which regulates the occurrence and development process of cell proliferation, apoptosis, differentiation and metabolism. Studies have shown that miRNA can affect the proliferation and differentiation of BMSCs by targeting and regulating the expression of related genes. However, whether microRNA can regulate psoralen-induced osteogenic differentiation and its specific mechanisms remain unclear. The aim of this study was to identify miRNA target genes regulating the osteogenic differentiation of BMSCs induced by psoralen and further analyze the mechanism.miRNA microarray was performed to screen the differentially expressed mirnas induced by psoralen, and bioinformatics analysis and luciferase reporter gene assay were used to determine the target sites of let-7i-5p. The mechanism of let-7i-5p in psoralen induced osteogenic differentiation of BMSCs was investigated using in vitro overexpression or inhibition approaches.The results showed that miRNA microarray analysis and reverse transcription quantitative PCR further confirmed that let-7i-5p was up-regulated during the process of psoralin-induced osteogenic differentiation of BMSCs.Bioinformatics analysis and luciferase reporter gene assay confirmed CKIP-1 as a potential target of let-7i-5p.Overexpression of CKIP-1 by transfection with CKIP-1 mimetic significantly inhibited the expression of BMP, Runx2, and Smad1, while CKIP-1 inhibitor promoted their expression.Overexpression of let-7i-5p partially relieved the inhibitory effect of CKIP-1 on osteogenic differentiation of BMSCs.Our results suggest that let-7i-5p regulates psoralen-induced osteogenic differentiation of BMSCs by targeting CKIP-1, which provides a new therapeutic target for osteoporosis.

## Introduction

Osteoporosis (OP), as a systemic metabolic bone disease, is characterized by low bone density and content, damaged physiological and anatomical structure of bone, and weakened strength of bone, etc[1]. It occurs in older women after menopause, clinical physical pain, low back pain with older women, mostly by osteoporotic fractures (osteoportic fracture, OPF), patients with severe osteoporosis is likely to see bone fracture delayed union or nonunion[2]. Estrogen can change the bone conversion rate by inhibiting bone resorption to maintain bone content. Postmenopausal women have degraded ovarian function, decreased estrogen level, enhanced bone resorption, bone balance imbalance, and bone mass loss, which are easy to cause osteoporosis[3]. Therefore, elderly women are prone to lumbar compression fractures, femoral neck fractures and other osteoporotic fractures after falls. Previous studies have shown that at least half of the global postmenopausal women are likely to suffer from osteoporosis[4]. The incidence of OP is positively correlated with age, and with the aggravation of aging population, its incidence will continue to rise. In today’s society, competition in all walks of life is fierce, people are under great pressure, working hours are significantly extended, time for rest and exercise is significantly reduced, physical fitness is generally decreased, the range of osteoporosis population is gradually expanded, and OPF will also increase exponentially[5]. OP has brought an increasing economic burden to the national medical insurance industry and individuals, and has become a disease of the global public health system[6].

Currently, the main treatment of OP is to promote bone formation and inhibit bone resorption[7], while estrogen replacement therapy (ERT), estrogen receptor modulators, basic supplements for bone health, calcitonin and bisphosphonates are also gradually used to reduce bone loss and thus prevent and treat osteoporosis. Teriparatide can improve the activity of osteoblasts and promote bone formation, but its high price limits its widespread application[8]. Studies[9] have reported that Raloxifene can reduce the incidence of lumbar compression fracture in elderly postmenopausal women, and has a good bone protection effect. However, long-term use of ERT may increase the risk of breast cancer and endometrial tumors in women, and bisphosphonates may induce adverse reactions such as mandibular osteonecrosis and atypical femoral fracture[10–11]. Estrogen receptor modulators are also associated with the risk of cardiovascular disease, venous thromboembolism, female massive uterine bleeding, and endometrial hyperplasia[12]. Therefore, in order to reduce these potential risks and further enhance the prevention and treatment of osteoporosis, Chinese herbal medicines with low side effects, high safety, strong efficacy and low price gradually become a research hotspot.

Psoralen is the main active ingredient extracted from the traditional Chinese medicine psoralen, which plays an important role in the treatment of bone injury diseases. Psoralen is recognized as a phytoestrogen, which can effectively stimulate the formation of new bone in local areas[13], inhibit osteoclast activity and reduce bone resorption[14], and regulate the relative balance between osteogenesis and osteoclast. Although many studies[15–19] have indicated that psoralen can promote the proliferation and differentiation of osteoblasts or inhibit osteoclasts by acting on TGF-β/Smad, BMP, MAPK, PI3K/AKT, Wnt/β-catenin, Hedgehog, Notch, OPG/RANKL, ER and other signaling pathways However, the specific molecular mechanisms, potential signaling pathways and targets of psoralen promoting bone formation and inhibiting bone resorption remain to be further elucidated and supplemented.

CKIP-1,a bone formation inhibitor, can work with ubiquitination regulatory factor 1(Smurf1) to regulate the breakdown of Runx and Smad proteins[20], playing an important role in the process of bone morphotransformation[21]. CKIP-1 can inhibit bone formation by interrupting the BMP/Smad signaling pathway, reducing bone mineralization and deposition. After silencing or knocking out CKIP-1 gene, osteoblast proliferation and differentiation are significantly active, and bone strength and bone density of mice are significantly improved[22]. It is suggested that we can target CKIP-1 small interfering RNA (CKIP-1 SiRNA) specific to osteoblasts to regulate the bone balance process. Taking CKIP-1, a negative regulator of bone formation, as the target breakthrough point, it is expected to find a new direction for the treatment of osteoporosis. MicroRNA (miRNA) is a small molecule single-strand non-coding RNA that regulates cell proliferation, apoptosis, differentiation, metabolism and other developmental processes[23–24]. Studies have shown that miRNA can affect the proliferation and differentiation of BMSCs by targeting the expression of related genes[25]. As a miRNA in the Let-7 family, let-7i-5p has been confirmed to have a close association with tumors, and may also play an important role in osteogenic differentiation of BMSCs. Studies[26] found that after overexpression of let-7i-5p, the expression of CKIP-1 was significantly reduced. Therefore, we believe that let-7i-5p may target and regulate the expression of CKIP-1, thus promoting osteogenic differentiation of BMSCs.

In this study, mouse bone marrow mesenchymal stem cells (mBMSCs) were treated with psoralen in vitro and in vivo to explore the influence of psoralen at different concentrations on osteogenic differentiation of mBMSCs, clarify the relationship between psoralen and let-7i-5p/CKIP-1 pathway, and clarify the molecular mechanism of psoralen’s influence on bone formation and bone resorption. In order to provide a new idea for anti-osteoporosis treatment.

## Materials and methods

### Materials

All C57BL/6 mice (24 females, 4 weeks old) were purchased from Huafukang Biotechnology Co., LTD. (Beijing,China), 4 of which were used as a reserve to prevent death. All mice were reared in SPF animal lab of Nanchang University. 293T cells were obtained from Beina Biology (Beijing,China), and mouse bone marrow mesenchymal stem cells were obtained from Seye Biology (Guangzhou,China). Psoralen (CAS 66-97-7) was purchased from Sourleaf Biology (Shanghai,China), estradiol and alkaline phosphatase staining solution was obtained from Solebaol (Beijing,China), alizarin red staining solution was purchased from Kaiji Biology (Jiangsu,China); All animal feeding and other experimental operations are conducted in the Animal Experimental Center of Nanchang University in accordance with the “Experimental Animal Guidelines” of the United States Institute of Health, in line with the relevant management guidelines and experimental animal ethical requirements, and obtained the approval of the Chinese Commission for Experimental Animals.

### Cell culture and identification

C57BL/6 female mice (120-140g) at the age of 4 weeks were selected and given peritoneal injection of 2% pentobarbital sodium 40mg/kg for anesthesia, then soaked and disinfected with 75% for 5 minutes. Both femurs were dissected on a super clean table, and the muscle was removed and cut in the middle of the femur. The alpha-MeM medium containing 10% fetal bovine serum was pumped out of the bone marrow from the broken end by a syringe. A single cell suspension with a density of 1×10^5^ cells/cm^2^ was prepared and inoculated in a 25cm^2^ culture flask, which was gently shaken to mix the cell density and placed in an incubator at 37 [and 5% CO_2_. After 48 hours, the culture medium was completely changed, and then the culture medium was changed every 3 days. After cell clones were gradually fused and spread to the bottom of the culture flask, the superneant of cell culture was discarded. The cells were washed twice with 1×PBS, and 0.25% pancreatin (containing 0.02% EDTA) was added for digestion.When the cells became round, the digestion was terminated by adding the medium, and cell suspension was collected into a 10mL centrifuge tube. The cells to be resuscitated were removed from liquid nitrogen and transferred into 39 [dry thermostatic heater via dry ice box; After the cells melted, they were quickly placed in a super-clean table, transferred to a centrifuge tube, and labeled for centrifugation (1000rpm,3min). The supernatant was discarded and fresh medium was added into the centrifuge tube to re-suspend the cells. The cell suspensions were divided into prepared cell culture bottles or petri dishes in a ratio of 1:3, and labeled and placed in the incubator. When the cells reached 80% fusion, they could be passed through. In this study, cells passed through to the third generation were used.

### Cell osteogenesis induction and psoralen treatment

The cells of the third generation were filled with the bottom of the culture bottle, and the supernatant of the cell culture was discarded, washed twice with 1×PBS, and digested by adding 0.25% pancreatic enzyme (containing 0.02% EDTA). When the cells became round, the digestion was terminated by adding the cell suspension into the 10mL centrifuge tube, centrifuged at 1000rpm for 3min, and the supernatant was discarded and added into the medium PBS to re-suspend the cells. Centrifuge, remove supernatant and repeat 3 times; The cell density was adjusted to 2×10^5^ cells /ml, and the culture medium was discarded. When the cells were well stuck to the wall and the cell density reached about 80%, the normal cell group, cell+psoralen group, cell+bone formation induction solution group (dexamethasone l0umol/L, vitamin C50umol/L, β-glycerol phosphate l0umol/L, α-MEM culture medium of 10% fetal bovine serum), cell+osteogenic induction medium+psoralen groups were treated and induced, respectively, and placed in a constant temperature incubator. The fluid was changed every 3 days for continuous induction for 14 days.

### miRNA microarray detection

The third-generation BMSCs were inoculated in 6-well plates at a density of 1x10^5^ cells /ml.After reaching 80%-85% confluence, BSMCs were induced by osteogenic induction solution and psoralfor 7 days (3 cases in each group) for miRNA microarray detection, and differentially expressed miRNAs were screened out. Reverse transcription quantitative PCR (RT-qPCR) was used to verify the differentially expressed mirnas.

### Reverse transcriptional quantitative (RT-q)PCR

RNA was extracted from the cells and tissues of each group, and cDNA was synthesized according to the reverse transcription kit. cDNA was used as the template for detection on a fluorescent quantitative PCR instrument, and β-actin was used as the internal reference. The following reagents were added in the following order: total volume 20μL. Nuclease free water 7.6μL, cDNA 2μL, Forward Primer 0.2μL, Reverse Primer 0.2μL, 2×All-in-One qPCR mix 10μL. The reaction conditions are as follows: pre-denaturation at 95°C for 10 minutes, then denaturation at 95°C for 40 cycles for 10 seconds, annealing at 52.3°C for 30 seconds, and finally extension at 72°C for 30 seconds. Then β-actin was used as the internal reference gene. The relative quantitative analysis was carried out by 2^-DDCt^ method. The relative expression levels of mmu-let-7i-5p, CKIP-1, Runx2, Smad1 and BMP2 in each group were calculated. The primer sequence is as follows:

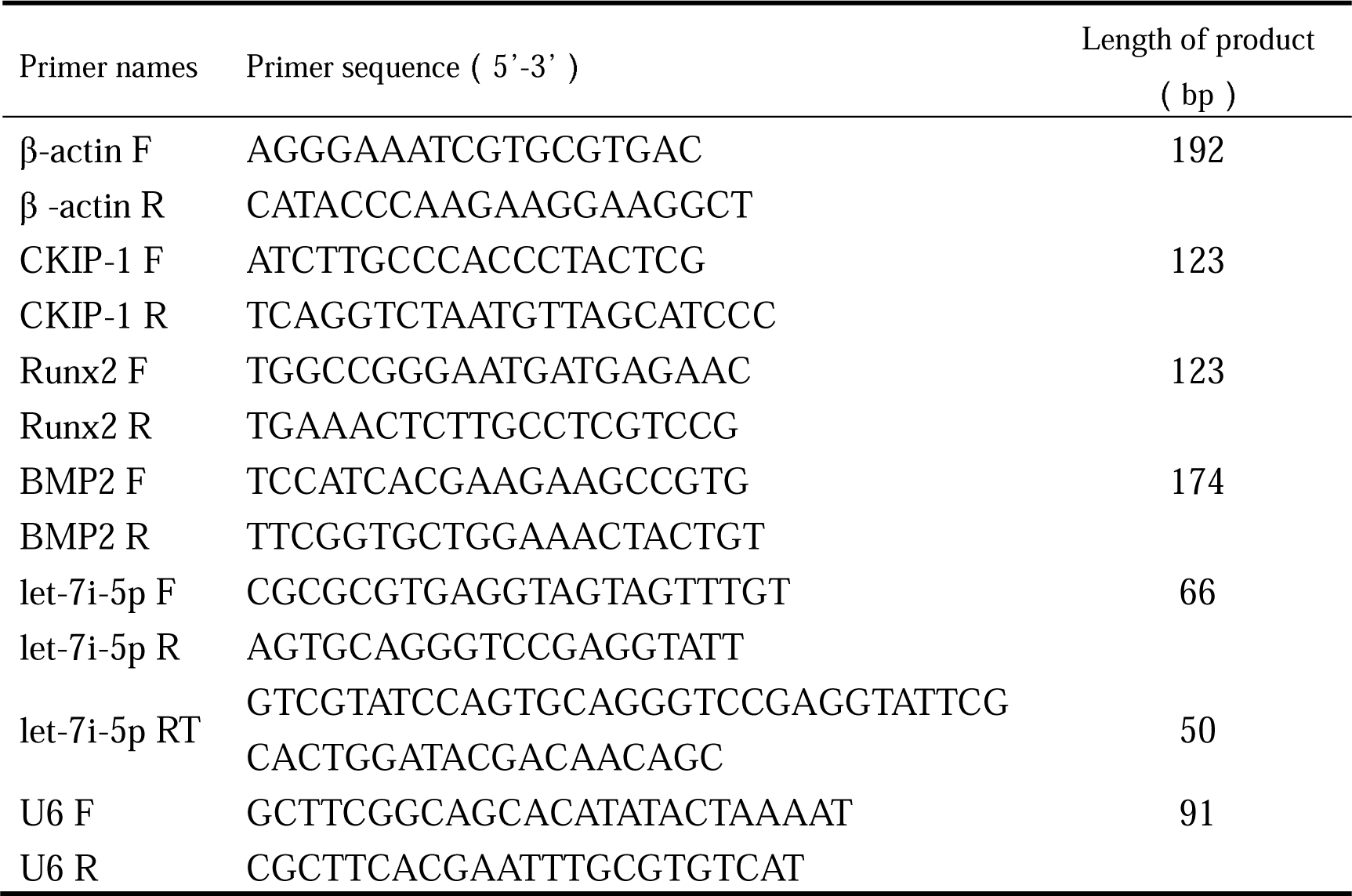

### Bioinformatics analysis

TargetScan (http://www.targetscan.org), NCBI (http://www.ncbi.nlm.nih.gov) and Ensembl (http://www.asia.ensembl.org) The potential let-7i-5p binding site was analyzed at the 3’-untranslated region (3’UTR) of CKIP-1.The consistency of the analysis and prediction of these three websites shows that they are reliable, and their possibility is further verified by in vitro experiments.

### Lentivirus packaging of overexpression vector and Virus Infection

HEK293T cells were cultured, and the lentivirus packaging experiment was started after the cell adhesion grew to about 70-90% density. The preheated 15mL complete medium was replaced 1h before the plasmid transfection experiment, and the Lenti-Mix-DNA transfection system was prepared. The preheated 20mL complete medium was replaced 8-10h after transfection, and the culture was continued. Fluorescence observation and photography were performed 24h after transfection. 72h after transfection, the supernitant of the cell culture medium was collected by filtering with a 0.45µM filter. The filtrated supernitant was fully mixed with 5×PEG8000 lentivirus concentrate and treated overnight at 4[. The centrifuged lentiviral particles were re-suspended with 1mL serum-free DMEM, then packaged and frozen at -80.

Fluorescence microscopy was used to detect the titer of lentivirus infection, and qPCR was used to absolutely quantify the number of lentivirus particles in the supernatant. After lentivirus infection was verified, the constructed plasmids were transformed, monoclonal expanded culture and deendotoxin extraction to obtain high quality endotoxin-free plasmids. Finally, a Lenti-Mix-DNA transfection system was prepared by mixing the target plasmid and no-load plasmid with Lenti-Mix, respectively. HEK293T cells were transfected together, lentivirus was packaged, and concentrated and frozen at -80 [after each tube.The supernatant of cell culture was discarded, washed twice with 1×PBS, and digested by adding 0.25% pancreatin (containing 0.02% EDTA). When the cells became round, the digestion was terminated by adding into the medium, and cell suspension was collected into a 10mL centrifuge tube, centrifuged at 1000rpm for 3min. The supernatant was discarded, then added into the medium and blown well with a pipette for counting. The above cell suspension was diluted to an appropriate cell density and then added into the prepared culture plate for culture at 37 [and 5% carbon dioxide incubator. Transfection was performed after the condition was good. After the cells were in good condition, the cell culture medium was changed into complete culture medium containing 5ug/ml staining aid polybrene. The number of cells in each well of the six-well plate was 2×10^5. According to the virus titer, lentivirus of corresponding volume was added to each well for transfection according to MOI (MOI=virus titer × virus volume/number of cells) of 30.16h after transfection, the fluid was changed, and 72h after transfection, downstream detection was performed for verification.

### Cell transfection

When the cell density reached 70%, it was ready for transfection. The cell medium was changed to serum-free medium with a volume of 1ml. Two sterilized EP tubes were taken, 125ul Opti-MEM was added to each tube, and 5uL lipofectamine 3000 was added to one tube. Another EP tube was added with 12.5ul let-7i-5p inhibitor (let-7i-5p inhibitor dry powder dissolved with DEPC water; 125ul/1OD), mixed and incubated at room temperature for 5min; The above two EP tubes were mixed well and incubated at room temperature for 15min. The mixed liquid drops were placed into the corresponding holes in the six-well plate, and the cells were put back into the incubator for culture. 4h after transfection, 1ml complete medium containing 20% serum was added into the six-well plate. The corresponding detection was carried out 48 hours later.

### Dual luciferase reporter gene detection

The experiment was divided into 7 groups with 3 multiple Wells in each group, and a total of 21 Wells of 293T cells were prepared in a 12-well plate. Group L (lipo3000) and Group P (P3000) were prepared. A group of 75ul was prepared from component P as a blank reagent group, and the rest group P was mixed with sea kidney plasmid. The P group containing the sea kidney plasmid was divided into 2 tubes with 225ul each, and the two groups were divided into 3 tubes with 75ul mixture each. After centrifugation, 125ul DEPC water was added into 1 OD (250ul DEPC water was added into 2OD) to prepare a 20uM storage solution (1000X) with a working concentration of 25nM. Add 1.25ul to each well. Each group was mixed with group L, adding 75ul to each tube. After mixing, incubate at room temperature for 15min, add 50ul cells to each well, add reagents to 12-well plates drop by drop, shake evenly after each group is added. Cells were collected 48h after transfection, and 200ul lysate was added to each well to split the cells. The cells were cleaved at 4 [for 20min, and the lysated cell samples were stored at -80[. Computer test: 70ul of cell lysate was taken from each group and added to the black 96-well plate, and 100ul of luciferase detection reagent was added to each well. Firefly luciferase activity was detected on computer. Then,100ul of cifluciferase detection reagent (detection substrate: buffer =1:100) was applied to each well to detect cifluciferase activity.

### Western blot detection

The cells and tissues of each group were added with lysate, lysis was performed on ice for 30min, centrifuged at 10000 rpm/min at 4 [for 10min, and the total protein was obtained by carefully drawing the supernatant. Protein concentration was determined by BCA kit. The protein was denatured, samples were taken, sodium dodecylbenzene sulfonate gel electrophoresis was performed for 1-2h, and the membrane was transferred wet for 30-50min. Primary antibody solution was incubated at 4 [overnight. Incubate the secondary antibody solution at room temperature for 1-2h. ECL exposure solution was dripped onto the film and exposed in a gel imaging system. Grayscale values of each antibody strip were analyzed using “Quantity one” software.

### Statistical analysis

SPSS19.0 software was used for statistical analysis. All experiments were repeated for 3 times, and quantitative results were expressed as mean±standard deviation (*X±S)*. Single factor analysis of variance was used for quantitative comparison among multiple groups, and LSD method was used for pound-for-pair comparison. Test level P=0.05, Graphpad7.0 was used for mapping, and Imagepro J software was used for gray value analysis.

## Results

### During the osteogenic differentiation of mBSMCs induced by psoralen, CKIP-1 expression was down-regulated, while let-7i-5p expression was significantly up-regulated

In order to study the different expression levels of miRNA during psoralen-induced osteogenic differentiation, BMSCs were treated with psoralen for 3 days, and then miRNA microarray detection was performed on the cells. A total of 91 mirnas were differentially expressed between the control group and the psoralen group. Compared with the control group, 54 mirnas were up-regulated and 37 mirnas were down-regulated in the psoralen group. The data is analyzed and illustrated in a heat map (Figure 1A). Among the miRNAs, 10 mirnas with the most significant differences in up-regulation were selected (let-7i-5p, miR-93-5p, miR-17-5p, miR-200c-3p, miR-96-5p, miR-92a-3p, miR-20a-5p, miR-148a-3p, LET-7I-5P).

**Figure 1.**
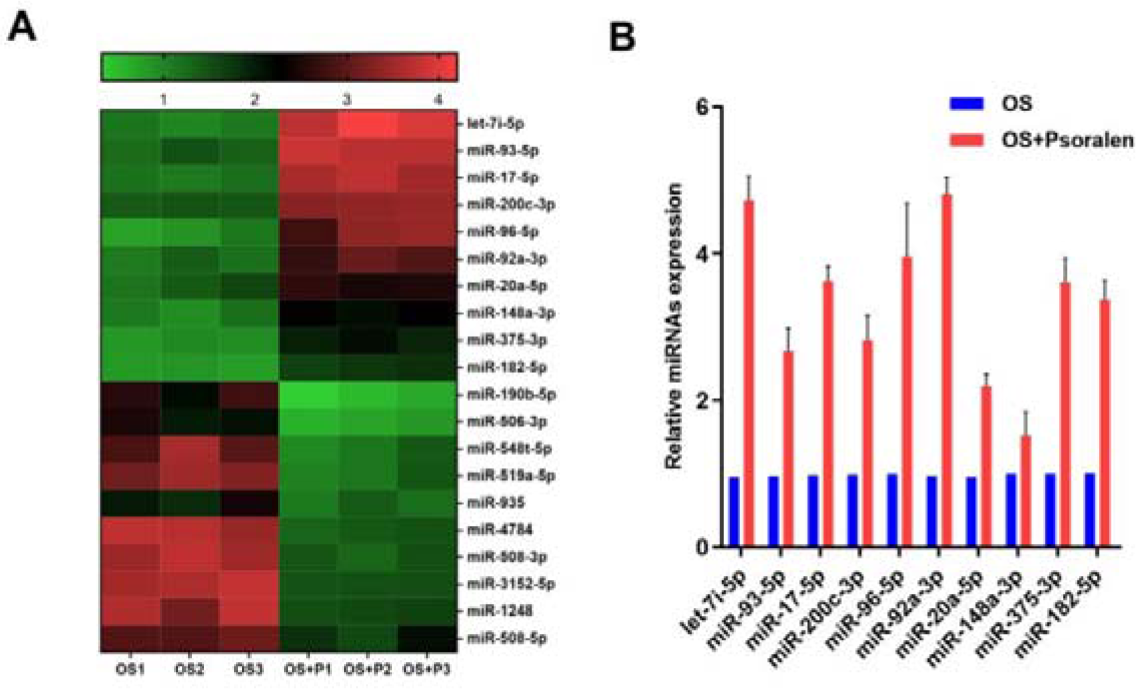
Heat map of miRNAs differentially expressed in bone marrow mesenchymal stem cells during osteogenic differentiation induced by psoralen. (A) Compared with the control group, 91 miRs were differentially expressed, of which 54 were up-regulated and 37 were down-regulated. Red means up, green means down. (B) The differentially expressed miRNAs were detected by qRT-PCR, and let-7i-5p expression was significantly up-regulated; OS:Osteogenic induction fluid; Psoralen:Psoralen induction fluid.

**Figure 2.**
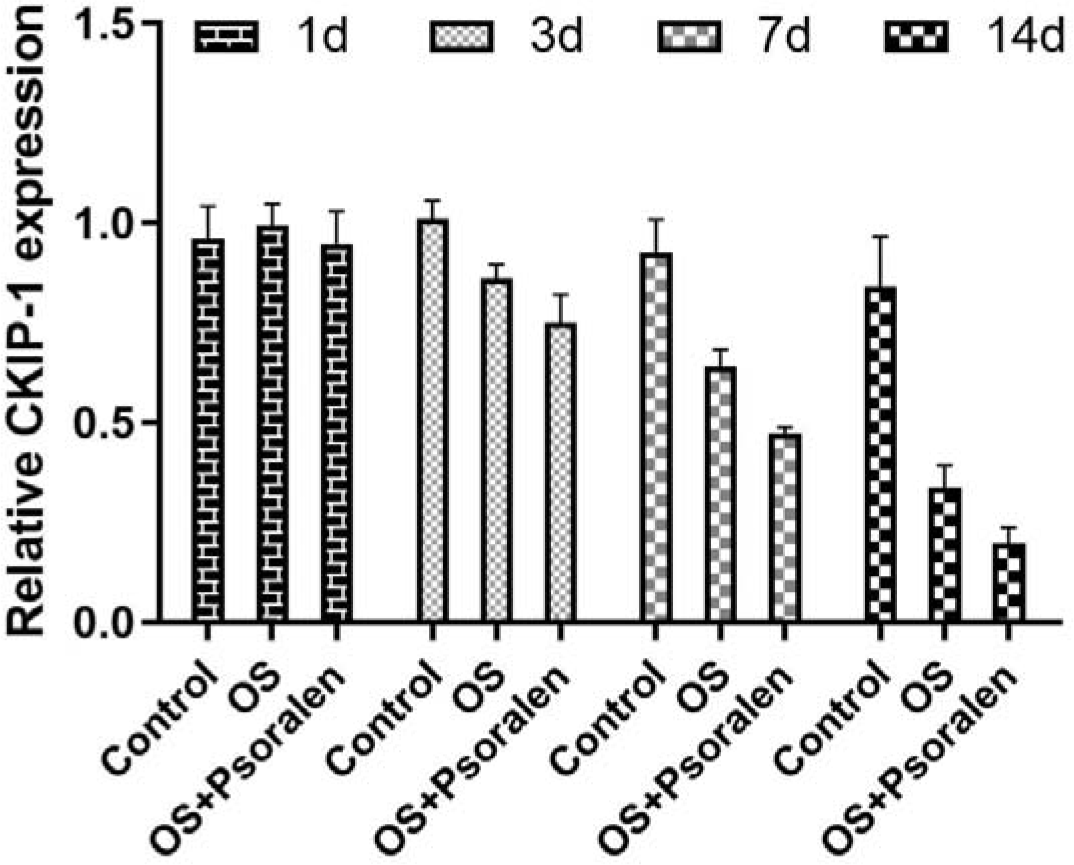
The expression of CKIP-1 in the osteogenesis of BMSCs induced by psoralen was detected by RT-PCR; OS:Osteogenic induction fluid;Psoralen:Psoralen induction fluid.

miR-375-3p, miR-182-5p) were validated to confirm the results of miRNA microarray analysis by RT-qPCR. Compared with the control group, the expression of all the above 10 miRNAs was up-regulated, among which let-7i-5p was the most significantly up-regulated (Figure 1B), which was basically consistent with the chip results. The third-generation BMSCs were cultured in groups and divided into blank group, osteogenic induction group and psoralen induction group. RT-PCR was used to detect the expression changes of CKIP-1 at different times of culture and induction. The results showed as follows: Compared with the blank group, the expression of CKIP-1 in the osteogenic induction group and the psoralen induction group was gradually down-regulated with the time of osteogenic induction, and the expression of CKIP-1 in the psoralen induction group was more significantly down-regulated than that in the osteogenic induction group. These results indicated that CKIP-1 expression was down-regulated during the osteogenic differentiation of BSMCs induced by psoralen.

### CKIP-1 overexpression vector and let-7i-5p inhibitor were successfully transfected

Western blot and qPCR were used to verify the transfection of mBMSCs with CKIP-1 overexpression vector and let-7I-5p inhibitor. The results were shown in Figure 5. let-7i-5p expression was significantly decreased in let-7i-5p inhibitor group (Figure 3A). Compared with control and no-loading, CKIP-1 mRNA and protein levels were significantly up-regulated in the CKIP-1 overexpression group (Figure 3B, 3D), indicating successful transfection of mBMSCs with CKIP-1 overexpression vector and let-7i-5p inhibitor.

**Figure 3.**
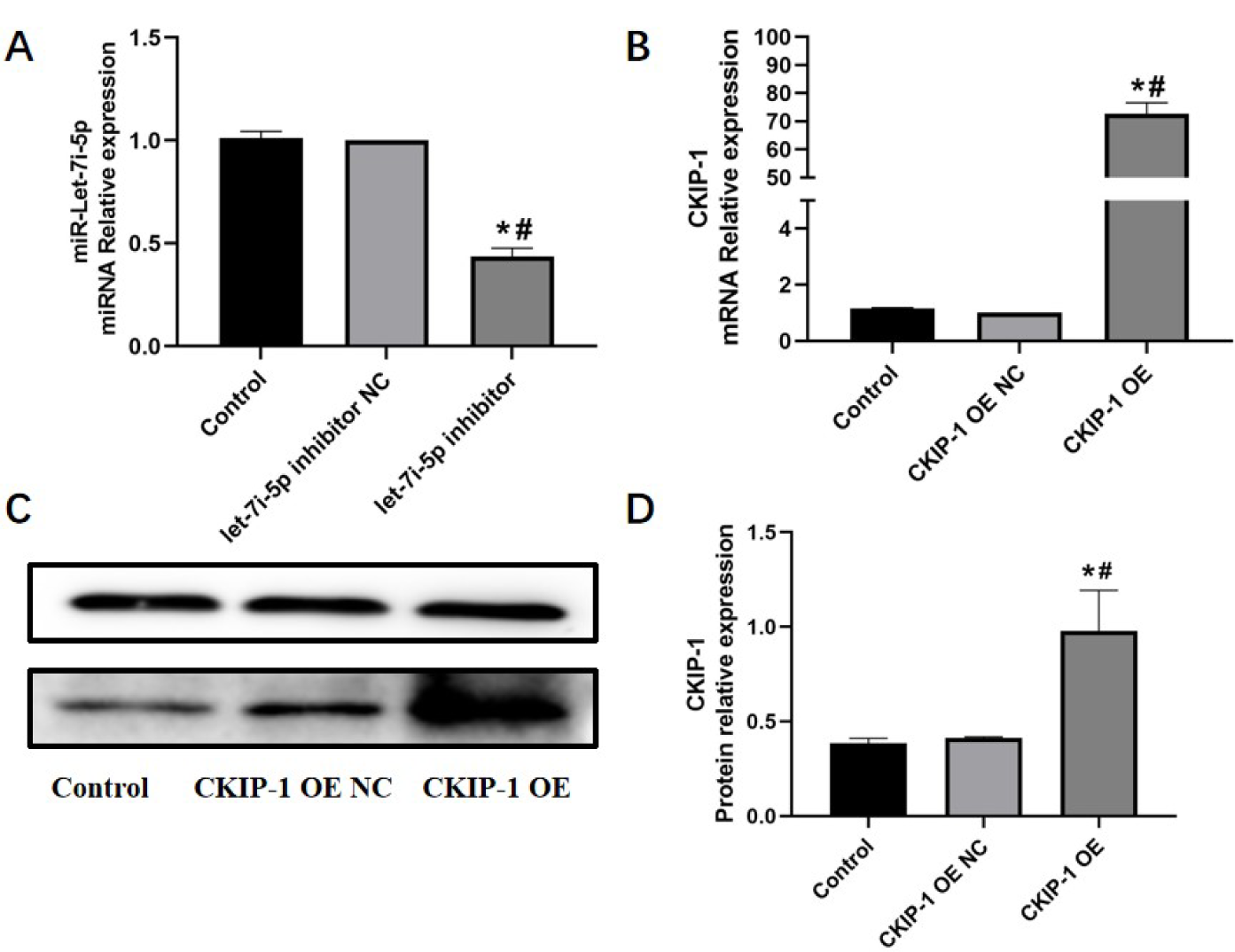
Western blot and qPCR for transfection verification (*P<0.05 VS Control; #P<0.05 VS CKIP-1 OE or let-7i-5p inhibitor NC; OE:overexpression; NC:normal control).

### CKIP-1 is a potential target binding site for Let-7i-5p

As shown in Figure 4, compared with the CKIP-1 WT, CKIP-1 mut and CKIP-1 WT+let-7i-5p mimic NC groups, the fluorescence value of 3 ’-UTR region in the CKIP-1 WT+let-7i-5p mimic group was significantly decreased.This indicates that there is a targeting relationship between let-7i-5p and CKIP-1, and CKIP-1 may be the target gene of let-7i-5p. To gain insight into the molecular mechanisms by which mirnas regulate osteogenic differentiation of BMSCs, potential mirnas were predicted using TargetScan and the transcription factor CKIP-1 was found to have a let-7i-5p binding site in its 3’UTR (Figure 4A). Therefore, further studies on let-7i-5p were conducted to determine whether let-7i-5p directly targeted CKIP-1 by luciferase reporter gene. The results showed that let-7i-5p mimics significantly inhibited the luciferase activity of wild-type CKIP-1 3’UTR. However, the luciferase activity of mutant CKIP-1 3’UTR was not ignificantly inhibited (Figure 4B).

**Figure 4.**
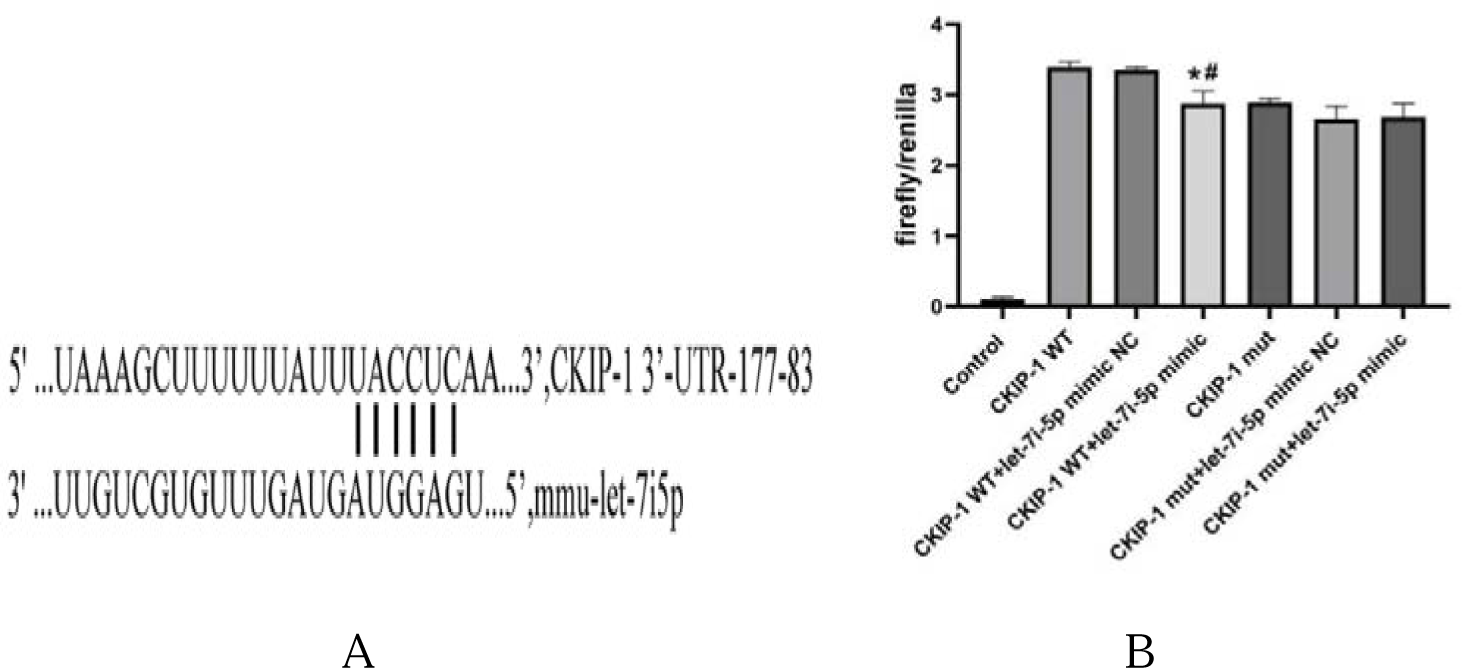
Targeting relationship verified by dual luciferase (*P<0.05 VS CKIP -1 WT; #P<0.05 VS CKIP-1 WT+let-7i-5p mimic NC,WT:wlid type; mut;mutan t type; NC:normal control;mimic:simulant).

**Figure 5.**
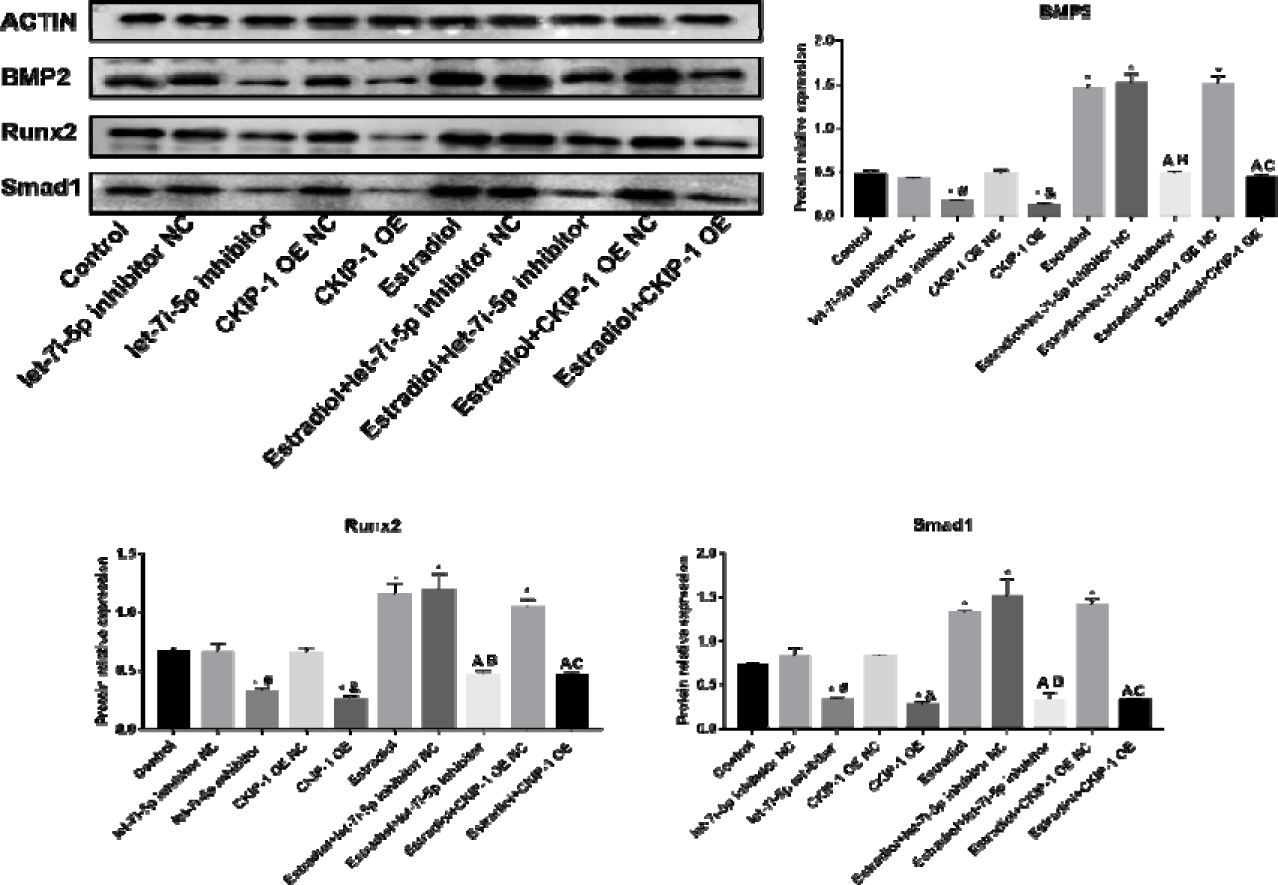
Western blot analysis of mBMSCs osteogenic differentiation related protein expression in each group (*P<0.05 VS Control; #P<0.05 VS let-7i-5p inhibitor NC; &P<0.05 VS CKIP-1 OE NC; A P<0.05 VS Estradiol; B P<0.05 VS Estradiol+let-7i-5p inhibitor NC; C P<0.05 VS Estradiol+ CKIP-1 OE NC;OE:overexpression; NC:normal control).

### CKIP-1 negatively regulates the osteogenic differentiation of BMSCs

In order to better understand the functional role of CKIP-1 in osteogenic differentiation of BMSC, the expression of Runx2, BMP and Smad1 was detected after BMSCs were transfected with CKIP-1 mimic and let-7i-5p inhibitor. As shown in Figure 5, mRNA and protein expression levels of Runx2, BMP, and Smad1 in the CKIP-1 overexpression group and the let-7i-5p inhibition group were significantly reduced compared with the control group and no-load group, indicating that CKIP-1 inhibited osteogenic differentiation of BMSCs. After adding estradiol to osteogenic induction, the degree of inhibition was relatively improved.As shown in Figure 6, compared with the control group and psoralatin induced group, the let-7i-5p inhibitor and CKIP-1 overexpression significantly inhibited the protein expression of BMP2, Runx2 and Smad1, indicating that CKIP-1 and let-7i-5p were involved in the regulation of the osteogenic differentiation of mBMSCs.

**Figure 6.**
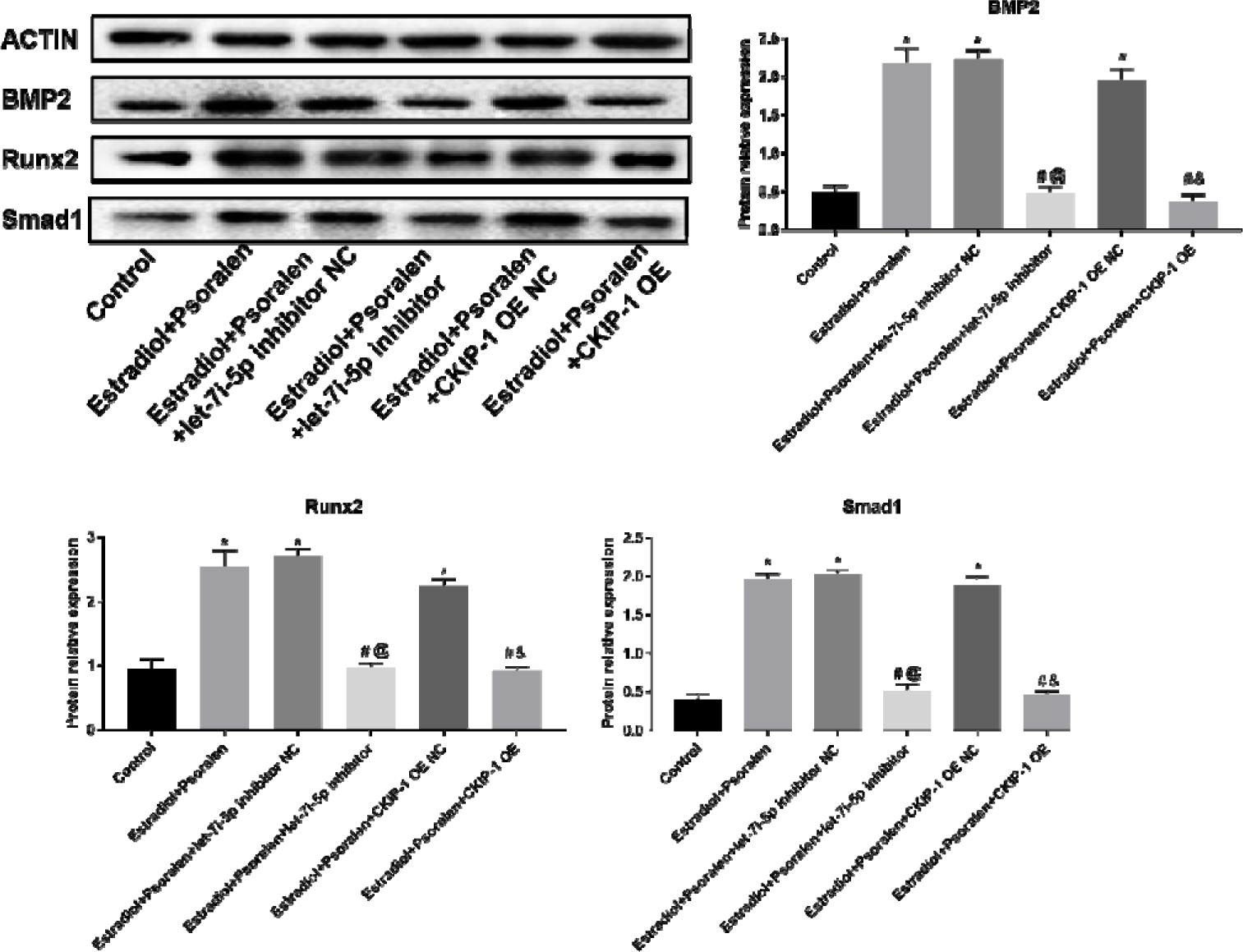
Western blot analysis of mBMSCs osteogenic differentiation related protein expression in each group (*P<0.05 VS Control; #P<0.05 VS Estradiol+Psoralen; @P<0.05 VS Estradiol+Psoralen+let-7i-5p inhibitor NC; &P<0.05 VS Estradiol+Psoralen+CKIP-1 OE NC; OE:overexpression; NC:normal control).

## Discussion

The maintenance of normal bone tissue depends on a dynamic balance mechanism formed between osteogenesis and osteoclasts. When bone balance is unbalanced, bone formation is reduced and bone resorption is increased, and osteoporosis is highly likely to occur. Related studies[27] reported that OPF, as the most common complication of osteoporosis, was found in nearly 20 cases of OPF worldwide within one minute. Therefore, osteoporosis patients need more safe, effective and accessible drugs to treat the disease and reduce the potential risk. Psoralen, as a furancoumarin-type phytoestrogen, has been regarded as an effective component of psoralen in the treatment of bone diseases. In addition to its known anti-inflammatory, anti-tumor and anti-cardiovascular functions[28], psoralen has also been confirmed to have the effect of promoting osteogenesis and inhibiting osteoclasts[29]. Bone marrow mesenchymal stem cells (BMSCs) can differentiate into osteoblasts, osteoclasts and adipocytes in multiple directions. Under certain stimuli, BMSCs can differentiate into osteoblasts with increased expression level of related osteogenic genes. Under psoralen treatment, small molecule RNA miR488 can increase the gene expression of osteogenic marker Runx2, promote the expression of ALP and osterix genes, promote the differentiation of BMSCs into osteoblasts, enhance bone calcium mineralization and deposition, and improve bone strength[30]. CKIP-1 negatively regulates bone formation, and miRNA let-7i-5p has been suggested to promote osteogenic differentiation of bone marrow mesenchymal stem cells. This study aims to further elucidate the relationship between let-7i-5p and CKIP-1 and the specific mechanism of psoralen promoting osteogenic differentiation of BMSCs.

In this study, the activity and proliferation of BMSCs were measured by CCK-8 method. It was found that psoralen promoted the proliferation of mBMSCs and maintained cell activity in a concentration-dependent manner. Alkaline phosphatase ALP is a bone conversion marker, which can not only be used to judge the osteogenic differentiation ability of BMSCs in the early stage, but also promote bone formation by improving the calcium deposition of BMSCs and the mineralization content of bone matrix[31]. Osterix has been confirmed as a transcription factor located in the downstream of Runx2 gene, which can cooperate with Runx2 gene to promote bone formation. After overexpression of Runx2, promoter activity in Osterix gene domain is enhanced, accelerating the differentiation and maturation process of osteoblasts[32]. Psoralen can not only reduce the bone resorption indirectly caused by rhBMP-2 in promoting osteogenesis, but also cooperate with rhBMP-2 to activate BMP signaling pathway and enhance signal transduction, thus improving the fracture healing rate of ovaries removed mice[33]. In order to further analyze the mechanism of psoralen promoting osteogenic differentiation of BMSCs, in this study, we targeted osteogenic genes such as BMP2, Runx2 and Smad1, and detected the mRNA and protein expression levels of these genes by fluorescence quantitative PCR and Western blot. These osteogenic markers have been previously confirmed to improve the activity of osteoblasts and promote cell proliferation and differentiation[34]. BMP2 belongs to the TGF-β family. As a key transcription factor, BMP2 can bind BMP receptors to activate the transcription of ALP, COLI and other genes, promote the proliferation and differentiation of osteoblasts, and accelerate bone formation[35–37]. Psoralen can improve the expression level of BMP-2, activate the transcription expression of downstream genes and promote the oriented osteogenic differentiation of mouse skull cells[15]. Runx2 is a specific transcription factor that promotes osteogenic differentiation of BMSCs by stimulating the expression of downstream proteins[38].

Certain concentration of psoralen can increase the expression levels of BMP, OPN, Runx2, Osterix and other osteogenic genes in hBMSCs and promote bone calcification deposition[19]. After Smad1 protein binds to receptors such as transcription factor BMP, its self-phosphorylation activates TGF-β/Smad signal transduction pathway and regulates the process of osteogenic differentiation of cells.

The osteogenic differentiation ability of BMSCs is significantly enhanced after treatment with low dose of psoralen[39].

miRNA let-7i-5p is involved in cell proliferation, differentiation, tumorigenesis and other signal transduction processes, regulates the transcription and expression of osteogenic genes, and plays an important role in promoting bone formation[40–41]. It is worth noting that studies[42] have also pointed out that let-7i-5p expression is reduced during osteogenic differentiation of hBMSCs, and after overexpression of let-7i-5p, the osteogenic ability of cells is reduced and bone formation is inhibited. CKIP-1 can participate in a variety of cell behavioral processes such as cell differentiation, apoptosis, and inhibition of bone formation, and play a role in regulating cell differentiation by virtue of its free shuttle function in the nucleus and cell membrane[43]. After CKIP-1 is combined with Smurf1, the degradation of Smad5 is accelerated, and the risk of OPF is significantly increased. After treatment with osteoblaster-specific small interfering RNA of CKIP-1, the damaged bone microstructure can be repaired[44]. In the process of BMSCs cultured in vitro, CKIP-1 silencing treatment significantly reduced the number of apoptosis, and increased the expressions of β-catenin and osteocalcin[45]. Therefore, drug target research design based on the interaction characteristics of CKIP-1 siRNA, CKIP-1 and Smurf1 is expected to become a new strategy for the treatment of osteoporosis.

In this study, microRNA agonists or miRNA inhibitors were used to promote or inhibit the expression of corresponding mirnas, while psoralen and estradiol were given intervention treatment to observe the expression of osteogenic genes and protein levels. During the experiment, we found that let-7i-5p could regulate the expression of CKIP-1, and the gene and protein expression of CKIP-1 decreased significantly after let-7i-5p was overexpressed. Combined with the results of double luciferase detection, we speculated that there might be corresponding binding sites between the 3 ’-UTR region of CKIP-1 and let-7i-5p. After CKIP-1 silencing, let-7i-5p expression was increased and osteoblast activity was significantly enhanced after psoralen treatment, which promoted bone formation.Therefore, we suggest that let-7i-5p targeting of CKIP-1 promotes osteogenic differentiation of bone marrow mesenchymal stem cells under psoralen.

However, this study still has the following limitations: First of all, only one concentration of psoralen was used in this experiment, and if multiple concentrations of psoralen were observed in control, the effect might be obvious. Secondly, the way in which psoralen promotes let-7i-5p to interact with its receptor still needs to be further explained. Finally. The specific upstream and downstream binding sites of psoralen involved in the let-7i-5p/CKIP-1 signaling pathway are unknown and need further study. In conclusion, psoralen may increase the expression level of let-7i-5p, inhibit the expression of CKIP-1, and increase the mRNA and protein levels of BMP2, Runx2 and Smad1, thus promoting bone marrow mesenchymal stem cell osteogenic differentiation, and thereby increasing bone strength and bone content, and finally achieving the purpose of treating osteoporosis.

## Acknowledgements

This research was supported by the National Natural Science Foundation of China (82060878).

## Availability of data and materials

The datasets used during the current study are available from the corresponding author on reasonable request.

## Ethical approval and consent to participate

Not applicable.

## Author’s contributions

CM, ZS and HS designed the research. ZXL, LH, and LGG performed the experiments. LHN,LB and YDM contributed to the statistical analysis of the data. CM,ZS and ZXL confirm the authenticity of all the raw data.ZXL and CM wrote the paper. All authors read and approved the final manuscript.

